# Amygdalo-nigral inputs target dopaminergic and GABAergic neurons in the primate: a view from dendrites and soma

**DOI:** 10.1101/2024.01.16.575910

**Authors:** JL Fudge, EA Kelly, TM Love

## Abstract

The central nucleus (CeN) of the amygdala is an important afferent to the DA system that mediates motivated learning. We previously found that CeN terminals in nonhuman primates primarily overlap the elongated lateral VTA (parabrachial pigmented nucleus, PBP, A10), and retrorubral field(A8) subregion. Here, we examined CeN afferent contacts on cell somata and proximal dendrites of DA and GABA neurons, and distal dendrites of each, using confocal and electron microscopy (EM) methods, respectively. At the soma/proximal dendrites, the proportion of TH+ and GAD1+ cells receiving at least one CeN afferent contact was surprisingly similar (TH = 0.55: GAD1=0.55 in PBP; TH = 0.56; GAD1 =0.51 in A8), with the vast majority of contacted TH+ and GAD1+ soma/proximal dendrites received 1-2 contacts. Similar numbers of tracer-labeled terminals also contacted TH-positive and GAD1-positive small dendrites and/or spines (39% of all contacted dendrites were either TH- or GAD1-labeled). Overall, axon terminals had more symmetric (putative inhibitory) axonal contacts with no difference in the relative distribution in the PBP versus A8, or onto TH+ versus GAD1+ dendrites/spines in either region. The striking uniformity in the amygdalonigral projection across the PBP-A8 terminal field suggests that neither neurotransmitter phenotype nor midbrain location dictates likelihood of a terminal contact. We discuss how this afferent uniformity can play out in recently discovered differences in DA:GABA cell densities between the PBP and A8, and affect specific outputs.

**Significance statement:** The amygdala’s central nucleus (CeN) channels salient cues to influence both appetitive and aversive responses via DA outputs. In higher species, the broad CeN terminal field overlaps the parabrachial pigmented nucleus (‘lateral A10’) and the retrorubral field (A8). We quantified terminal contacts in each region on DA and GABAergic soma/proximal dendrites and small distal dendrites. There was striking uniformity in contacts on DA and GABAergic cells, regardless of soma and dendritic compartment, in both regions. Most contacts were symmetric (putative inhibitory) with little change in the ratio of inhibitory to excitatory contacts by region.

We conclude that post-synaptic shifts in DA-GABA ratios are key to understanding how these relatively uniform inputs can produce diverse effects on outputs.

## INTRODUCTION

Across species, the dopamine system comprises three well-known subregions: ventral tegmental area (VTA, A10), the substantia nigra, pars compacta (A9) and the retrorubral field (A8)(Dahlstrom and Fuxe, 1964). In each area, DA neurons are embedded among GABAergic neurons (Oertel et al., 1982). Although the overall ratio of DA to GABAergic neurons is approximately 3:1, the ratios of DA to GABAergic neurons varies widely across the subregions, including in nonhuman primates (Nair-Roberts et al., 2008; Kelly et al., 2022). These shifting ratios could be a read-out of the GABAergic neurons’ relative propensity to participate in long-range projections, or their capacity to be intrinsic inhibitors of DA neurons via axon collaterals(Carr and Sesack, 2000; Tepper and Lee, 2007; Omelchenko and Sesack, 2009).

The midbrain DA neurons themselves are known to be connectionally and electrophysiologically diverse along medial-lateral and rostrocaudal organization of the subregions (Haber et al., 2000; Matsumoto and Hikosaka, 2009; Lammel et al., 2011; Hnasko et al., 2012; Mingote et al., 2015; Beier et al., 2019). The A10 group, or VTA, is the collective name for several distinct DA neuron subgroups, which were classically considered part of the ‘mesolimbic’ path, mediating reward learning (Berridge and Robinson, 1998; Schultz, 1998). However, different connections and functions of the A10 group are now associated with its medial and lateral aspects. Moreover, glutamatergic neurons, largely confined to the midline VTA, also participate in reward learning (Root et al., 2016). In contrast to the ‘midline’ VTA subnuclei, the laterally placed parabrachial pigmented nucleus (PBP) appears to have different inputs/outs and functions(Haber et al., 2000; Fudge et al., 2017; de Jong et al., 2019). The PBP accounts for an especially large area of the A10 in higher species (Halliday and Tork, 1986; McRitchie et al., 1996). Stretching from under the red nucleus and over the entire A9 region (pars compacta), the PBP A10 neurons meet the A8 neurons in the lateral midbrain. The A8 group extends caudolaterally to become dispersed among fibers of the medial longitudinal fasciculus(Jimenez-Castellanos and Graybiel, 1987; Gaspar et al., 1993; Francois et al., 1999). A8 neurons project to the striatum, temporal lobe (entorhinal cortex and amygdala(Cho and Fudge, 2010))(Deutch et al., 1988) and in primates, to the prefrontal cortex (Gaspar et al., 1992; Williams and Goldman-Rakic, 1993). Interestingly both the A10 and the A8 DA neurons share high levels of the calcium binding protein (CaBP), distinguishing them from the A9 neurons which lie beneath.

The central nucleus (CeN) of the amygdala is the major conduit between the main amygdala nuclei and the DA system. The amygdalonigral path is involved in both appetitive and aversive learning and behaviors (Price and Amaral, 1981; Gonzales and Chesselet, 1990; Zahm et al., 1999; Fudge et al., 2017; Steinberg et al., 2020). At least a subset of neurons in the amygdalonigral path co-contain corticotropin releasing factor (CRF) (Rodaros et al., 2007; Fudge et al., 2017; Steinberg et al., 2020). Importantly, the CeN projection to the ventral midbrain largely avoids the midline subnuclei of the A10 (midline VTA) in mice, rats, and monkeys (Deutch et al., 1988; Gonzales and Chesselet, 1990; Wallace et al., 1992; Fudge and Haber, 2000; Lee et al., 2005; Steinberg et al., 2020), terminating instead over the elongated parabrachial pigmented nucleus (PBP, or ‘lateral VTA’) and also the retrorubral field (A8) (Fudge and Haber, 2000; Fudge et al., 2017). In mice, general analyses of afferents to the ventral midbrain show a lack of biases for DA versus GABA cell types (Beier et al., 2015; Faget et al., 2016; Beier et al., 2019). Almost nothing is known of these relationships outside the midline VTA, or in higher species. A basic question is whether the CeN-nigral path terminates onto DA neurons, GABAergic neurons, or both.

An important issue in studying afferents onto DA and GABAergic neurons is that terminals have differential effects depending on relative contact in dendritic compartments and proximity to the axon. In both DA and GABAergic neurons, afferent terminal contacts near the soma and large proximal dendrites are relatively physiologically potent (Hausser et al., 1995; Seutin and Engel, 2010; Moubarak et al., 2022). This is due their proximity to the axon initial segment, which integrates incoming electrical activity and ultimately produces action potentials (Grace and Bunney, 1983; Tepper et al., 1987; Meza et al., 2018). Yet, the majority of afferent terminals are onto the vast network of small, non-proximal dendrites, which form the neuropil. DA neurons are known for extremely long dendrites with modest branching, while GABAergic neurons have highly complex dendritic trees (Oertel and Mugnaini, 1984; Tepper and Lee, 2007). To consider how the amygdalo-nigral path contacts these dendritic compartments in DA and GABAergic neurons, we used two methods biased towards visualization of terminals on 1) proximal dendrites and soma and 2) more distal dendritic sites, respectively. We first placed anterograde tracer injections into the CeN and created a mesoscopic map of the projections.

We then used confocal microscopy analyses to isolate and quantify tracer-labeled contacts onto tyrosine hydroxylases (TH) positive and glutamate dehyroxygenase-1 (GAD1) positive neuronal soma/proximal dendrites. We then used electron microscopy approaches to quantify tracer-labeled terminals onto smaller dendrites labeled TH- and GAD1-immunoreactivity. For the latter analyses, the proportions of inhibitory-versus excitatory-type contacts on each neuronal type, in the PBP and A8 regions of the terminal field, were assessed.

## MATERIALS AND METHODS

### Design

A total of 5 adolescent macaques (4 *m. fascicularis* and 1 *m.nemestrina*) (3 females, 2 males) ranging from 1.4-4.2 years of age received injections into the central nucleus (World Wide Primates, Tallahassee, FL, USA) (**Table 1**). After a survival period of 2 weeks, brains were harvested and processed to facilitate a step-wise process of analysis from the mesocopic to microscopic and ultrastructural levels (**Fig. 1**, **Table 1**).

**Figure 1.**
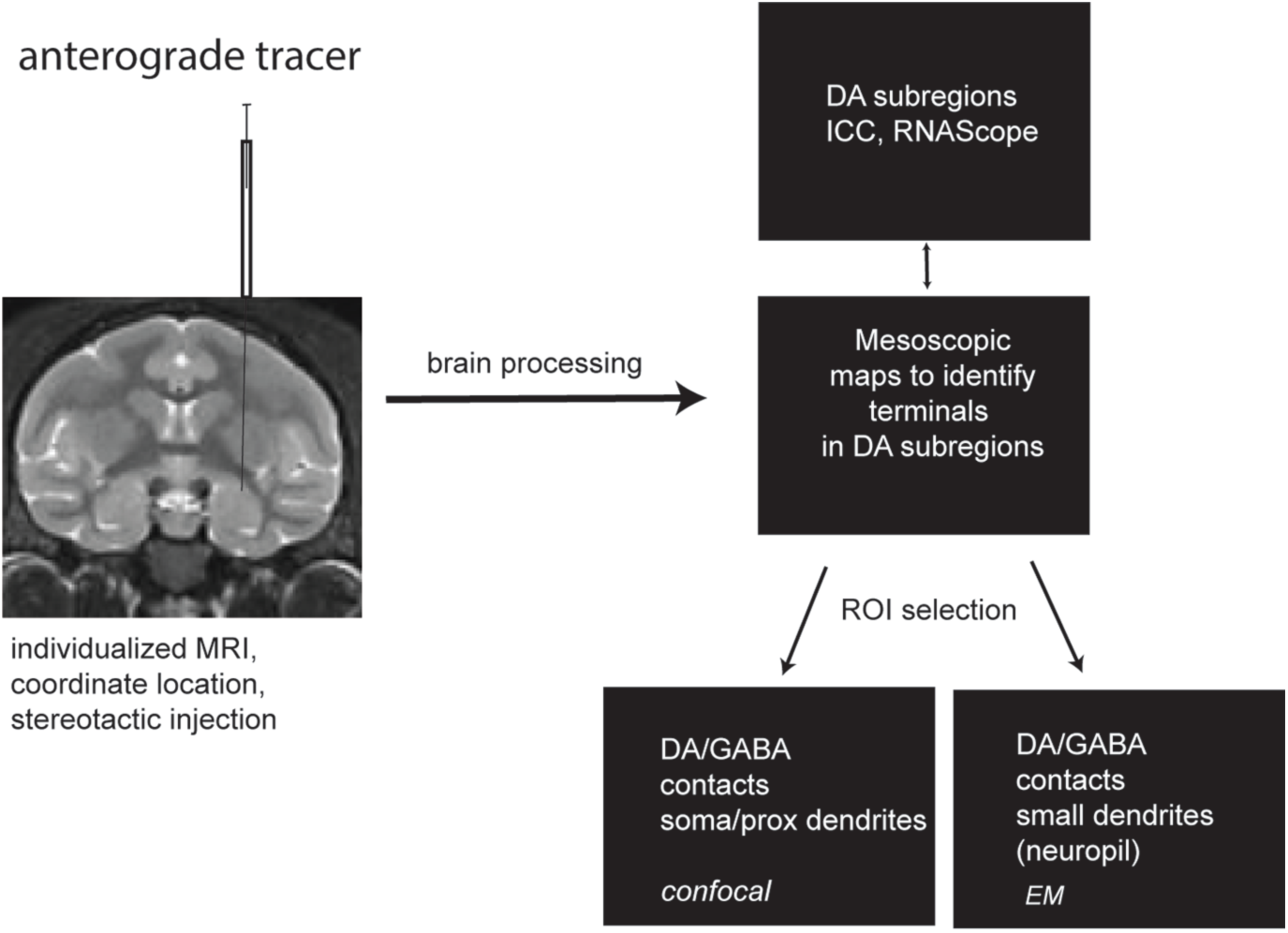
Design of experiments. Central nucleus was injected in five animals under stereotaxic guidance. Brains were processed to enable assessment of anterograde tracer expression at several microscopic levels.

### Surgical techniques

Experimental design and techniques were aimed at minimizing animal use and suffering and were reviewed by the University of Rochester Committee on Animal Research. To minimize animal use, several of these injections were performed as parts of other studies. Stereotaxic coordinates were determined for each animal prior to injection using individualized MRI imaging (3Telsa, coronal sections, 0.8 mm thick, 0.1 mm apart). For surgery, initial anesthesia was administered by intramuscular injections of ketamine hydrochloride (10mg/kg). Animals were intubated, and a deep plane of anesthesia was induced by isofluorane gas, maintained as needed during surgery. A craniotomy was performed, and the bone flap removed to visualize cortical surface landmarks. Small deposits of tracer were pressure-injected over 10-15 min into the central nucleus with a 0.5 μL Hamilton syringe with an attached glass pipette (tip diameter at 40 μm) (Hamilton Company, Reno, NV). The syringe was left in place for an additional 20 minutes to prevent leakage of tracer up the needle track. Only one injection for each type of tracer into the central nucleus was used per animal. Following injection placement, the bone flap was replaced, and overlying muscles and skin sutured. Two weeks after surgery, animals were deeply anesthetized and killed by perfusion through the heart with 0.9% saline containing 0.5 ml of heparin sulfate (200 ml/min for 10 minutes), followed by cold 4% paraformaldehyde (PFA) in a 0.1 M phosphate buffer/30% sucrose solution (100 ml/min for 1 h). The brain was extracted from the skull, placed in 4 % PFA overnight, and then put through increasing gradients of sucrose (10%, 20%, 30%) until the brain sunk. Brains were cut on a freezing microtome (40 μm) and all sections were stored in cryoprotectant solution (30% ethylene glycol and 30% sucrose in 0.1 M phosphate buffer) at -20 °C (Rosene et al., 1986) for use in various stages of the experiments moving from mesoscopic analysis and region of interest selection at the level of light microscopy through higher resolution imaging (**Fig 1**, **Table 1**).

### RNAScope processing

Tissue from one adolescent male animal (MF38, 3.3 years) used in our prior stereological studies was assessed for the distribution of TH and GAD1 transcripts in the DA subregions. Representative sections through the ventral midbrain through the rostrocentral and caudal midbrain were processed for TH and GAD1 mRNA. Sections from perfusion-fixed, cryoprotected sections (described above, 40 um) were thoroughly rinsed overnight in 0.01M phosphate buffered saline (PBS). The midbrain was blocked, excised and mounted out of buffer onto superfrost plus slides and stored up to 6 months at -80° until ready for processing.

Target probes were designed for a 2-plex fluorescent assay for macaque (Mmu) as follows: Mmu-TH (tyrosine hydroxylase [TH] transcript variant X1); Mmu-GAD1 and glutamate decarboxylase (GAD1)(Advanced Cell Diagnostics, ACD Bio, Newark CA). Multiplex RNAScope experiments are processed using RNAse-free reagents on-slide using RNAScope detections kits and reagents following the manufacturer’s protocol with slight modification to improve adhesion of thicker tissue (Biancardi et al., 2021). Sections were allowed to thaw and air dry, then washed for 10 min in double distilled water (ddH_2_O). Sections were postfixed for 30 min at 4°C in 4% paraformaldehyde (PFA), rinse 2 times in 0.01M PBS and then gradually dehydrated in ethanol (50%, 70%, and 2X 100% for 5 min each). Slides were baked in a HybEZ oven (in humidity control tray without humidifying paper) for 20 min at 60°C. Sections were treated with 1% H_2_O_2_, rinsed briefly in ddH_2_O and baked again for 10 min at 60°C. Remaining steps were as described in the ACDBio RNAScope protocol (Document # 323100-USM).

Following pretreatment, slides were air dried and incubated in protease III for 30 min at 40°C in HybEZ oven. Slides were then rinsed in distilled water and incubated in a 1-part C2/C3: 50-part C1 target probe concentration for 2 hours in HybEZ oven at 40°C. Tissue on slides was then processed using the RNAScope Multiplex Fluorescent Assay kit version 2 (ACD Bio) according to manufacturer’s instructions. The following Opal Dyes were used: Opal 520 reagent pack (FP1487001KT; Akoya Biosciences), Opal 570 reagent pack (FP1488001KT). Following Opal dye application, Trueblack (Biotium, Cat # 23007) was used at 1:20K dilution in 70% ETOH for 45 sec at room temperature to quench lipofuscin autofluorescence. Sections were washed in 1X wash buffer (ACD Bio) and coverslipped using Prolong Gold anti-fade mounting media (Invitrogen).

### Immunocytochemistry (ICC) processing

The antibodies used for detection of the specific tracer molecules have been documented in numerous tract-tracing studies including our own, and the distribution of anterogradely labeled fibers in the ventral midbrain was similar to previous results in a different cohort of animals (Fudge and Haber, 2000; Lanciego and Wouterlood, 2011). In the ventral midbrain, the pattern of calbindin-D28K (CaBP) and tyrosine hydroxylase (TH) immunoreactivity (IR) matched that found in monkey and human (Lavoie and Parent, 1991; McRitchie et al., 1996; Kelly et al., 2022). To identify the relative location of the CeN injection sites, we used cresyl violet and acetylcholinesterase (AChE) staining and corticotropin releasing factor (CRF)-IR, and somatostatin (ST)-IR in adjacent sections to distinguish the central nucleus (Heimer et al., 1999; Kovner et al., 2019; Fudge et al., 2022).

For visualization of tracers, 1:8 sections through the brain were rinsed in 0.1 M phosphate buffer with 0.03% Triton X (PB-TX), preincubated in 10% normal goat serum diluted in PB-TX (NGS-PB-TX), and then in placed in solutions with primary antibodies FR (1:1000, Invitrogen #A6397, made in rabbit), FS (1:500, Invitrogen, #A11095, made in rabbit) or LY(1:1000, Invitrogen, #A5750, made in rabbit) antibodies, and incubated for 96 h at 4°C. Sections were then rinsed 6 x 15 minutes in 0.1MPB-TX, blocked with 10% NGS-TX, and incubated with secondary biotinylated rabbit antibody (1:200). Following rinsing, the molecules were visualized using avidin-biotin reaction, colorized with 3’, 3’ diaminobenzidine.

The procedure for labeling adjacent sections for TH and CaBP-IR (ventral midbrain) and CRF and SST-IR (basal forebrain) was similar, using permanent labeling techniques. Adjacent compartments through the ventral midbrain were processed using mouse anti-CaBP (1: 10,000, C9848, Sigma, St. Louis, MO) and mouse anti-TH (1:10,000, MAB318, EMD Millipore Corp, Temecula, CA). Sections adjacent to tracer-labeled compartments through the basal forebrain were immunoreacted for CRF (BMA Biomedicals, previously Peninsula Laboratories, Switzerland, #T-4037, 1:5000, rabbit) and SST (Immunostar, #20067, 1:1000, rabbit) peptides. AChE staining was performed using the Geneser technique (Geneser-Jensen and Blackstad, 1971).

### Double immunofluorescent labeling for tracer/TH and tracer/GAD1 for confocal studies

To determine the relationship of anterogradely labeled fibers from CeN onto DA versus GABAergic cells in in the ventral midbrain, we processed adjacent compartments through the midbrain for double immunofluorescent staining for tracer and TH-IR, and tracer and GAD1-IR. Optimization and specificity of fluorescent staining for all antigens was first conducted in single labeling experiments, with reference to the permanently labeled compartments for tracer (above). Sections were rinsed 4 x 15 minutes in PB-TX and then rinsed on a rocker overnight at 4°F. The following day, tissue was then thoroughly rinsed with PB-TX and placed in a blocking solution of 10% normal donkey serum in PB-TX (NDS-PB-TX) for 1 hour. Following rinses in PB-TX, tissue was incubated in 3% NDS-PB-TX primary antisera to the tracer--either FR (1:1000, Invitrogen #A6397, made in rabbit), FS (1:500, Invitrogen, #A11095, made in rabbit) or LY (1:500, #A5750, made in rabbit)-- and anti-sera to TH (1:10,000, MAB318, EMD Millipore Corp, Temecula, CA) or GAD1 (1:10,000, MAB5406, EMD Millipore Corp, Temecula, CA, made in mouse) for ∼96 hours at 4°C. Tissues were then rinsed with PB-TX, blocked with 10 % NDS-PB-TX and incubated in the dark in secondary fluorescently tagged anti-mouse antibody for either TH or GAD1 (1:200, MAB5406, EMD Milipore)) and fluorescently tagged anti-rabbit antibody for tracer (1:200, AlexaFluor, Invitrogen), overnight at 4°C. Tissue was mounted out of 0.1M PB, pH 7.2 and cover-slipped with Prolong Gold anti-fade mounting media (Invitrogen).

### EM double immunolabeling for tracer/TH and tracer/GAD1

For EM studies, we chose 2 cases with FR injections into the CeN (one male and one female), to control for tracer (MNJ1 and MFJ55). One compartment for FR/TH and FR/GAD1 was selected from each animal for double immunoperoxidase (FR) and immunogold/silver labeling (TH or GAD1). Sections were dually stained for tracer-IR and either TH or GAD-IR. Briefly, sections were washed thoroughly in freshly-made filtered 0.05M phosphate buffered saline (PBS), subjected to 0.1% sodium borohydride in 0.1M filtered 0.1M PB to block active aldehydes (Craig, 1974), and blocked in a solution containing 3% NGS and 1% bovine serum albumin (BSA). Sections were incubated for 72-hours in the same blocking solution with anti-FR (1:500, Invitrogen #A6397, made in rabbit) and either anti-TH (1:5K, MAB318, EMD Millipore Corp, Temecula, CA, made in mouse) or anti GAD1 (1:1K, MAB5406, EMD Millipore Corp, made in mouse) at 4°C on a shaker followed by 24-hours at room temperature (RT). Sections were then washed in filtered 0.05M PBS and incubated for 1 hour in goat anti-rabbit IgG conjugated to biotin (1:200, Jackson ImmunoResearch, BA-1000), followed by ABC (per product instructions; Elite ABC Kit, Vector Labs, PK-6100). Tracer immunoreactivity was visualized with 3,3’-diaminobenzidine (0.5mg/ml; DAB) and hydrogen peroxide (0.03%) in 0.1M PB. Sections were rinsed thoroughly in filtered 0.05M PBS, followed by a second pre-incubation in a washing buffer solution containing 3% NGS, 0.8% BSA, 0.1% cold water fish gelatin in 0.1M PBS.

Sections were then incubated for 24 hours in goat-anti-mouse immunogold (Nanogold, N24915, Nanoprobes Inc., Yaphank, NY) in washing buffer at RT on a shaker. Following a thorough wash, tissue was incubated in 2.5% glutaraldehyde for 10 minutes at RT. Sections were rinsed thoroughly in 0.1M PBS, followed by a series of washes in Enhancement Conditioning Solution (1X concentration per package instructions, Electron Microscopy Science (EMS), Cat 25830, Hatfield PA). Sections were silver enhanced using the Aurion R-Gent SE-EM Kit (EMS, Cat #25520-90). Following additional post-fixation in 1% osmium tetroxide (EMS, Cat #19150), sections were dehydrated in ascending concentrations of ethanol (50%, 70%, 80%, 95%, 100% x2). Contrast was enhanced using 1% uranyl acetate in 70% ETOH within the dehydration series (EMS, Cat 22400). Sections were then treated with propylene oxide, impregnated in resin (Embed 812, Araldite 502, DDSA, DMP-30 (all EMS)) overnight at RT, mounted between ACLAR embedding films (EMS, Cat# 50425) and cured at 55°C for 72-hours.

Using immunolabeled near-adjacent sections, photomicrographs and digitized traces were used as reference to isolate tracer fiber innervation in PBP and A8 regions. Regions of interest were excised in a trapezoid shape (to record anatomical orientation) from the embedding films and re-embedded at the tip of resin blocks. Ultrathin sections (60-80 nm; evidenced by the sections silver sheen) were cut with an ultramicrotome (Reichert Ultracut E) and collected in sequence on bare square-mesh nickel grids (EMS, G200-Ni).

### Image Acquisition and Analyses

#### Light microscopy maps

The distribution of labeled fibers in the ventral midbrain was manually charted with darkfield microscopy under 10x illumination. Under lower power, the outlines of the surrounding structures, fiber tracts, and blood vessel were also mapped. Paper charts were then scanned into digital images and placed into Adobe Illustrator files. In adjacent sections, the boundaries of CaBP-positive neurons, CaBP-negative neurons, and the pars reticulata were mapped from adjacent sections using automated software (Neurolucida, Microbrightfield). The boundaries of the DA neurons and their dendritic fields were similarly drawn with the 4x objective, including major fiber tracts, third nerve fascicles, and blood vessels. The Neurolucida images were exported as SVG files and then imported into Adobe Illustrator software where the drawings for each section were scaled, and carefully overlaid onto scans of labeled fibers, matching fiducial markers. Each injection site was analyzed with respect to the location of labeled fibers in specific DA subregions based on established criteria (Kelly et al., 2022).

#### TH and GAD1 mRNA cellular distribution in DA subregions with RNAScope

Images adjacent to sections immunoreacted with permanent colorant (DAB) were used to validate and align 2-plex RNA transcript labeling. Sections in the rostral and caudal midbrain were assessed for TH- and GAD1-mRNA transcript in sequential channels on a Nikon A1R HD Laser Scanning Confocal microscope with NIS-Elements (Center for Advanced Microscopy and Nanoscopy) software.

The following excitation lasers (ex) and emission filters (em) were used for each label: GAD1: 520; ex 488, em 525/50 and TH: Opal 570; ex 561, em 595/50. After optimization, the illumination parameters for each channel were held constant for all sections. A tiled ROI of the ventral midbrain was then collected using a 10x/0.4NA Plan Apochromat objective using the lower magnified image as reference. Tiled images were visualized in Photoshop without channel corrections, cropped, and the boundaries of the subregions were mapped by aligning landmarks from CaBP-immunoreacted adjacent sections, with reference to fiducial markers such as fiber tracts and blood vessels.

#### Confocal Microscopy: Tracer contacts on TH- and GAD1-IR soma and proximal dendrites

Based on projection field maps showing that the amygdalo-nigral labeled terminals mainly encompassed the PBP and A8 subregions, these were selected for analysis in each case. Dual immunofluorescent images were collected on a Nikon A1R HD Laser Scanning Confocal with NIS-Elements (Center for Advanced Light Microscopy and Nanoscopy) software using tissue landmarks on adjacent projection maps. The following excitation lasers (ex) and emission lasers (em) were used: AlexaFlour 488; ex 488, em 525/50 and AlexaFluor 568; ex 561, em595/50.

Overview images were collected using a 20x/0.75NA Nikon Plan Apochromat VC objective to located ROIs in regions of tracer labeled fibers, and z-series stacks were selected and collected using a 60x/1.49NA Nikon Apochromat TIRF objective (xy pixel size0.17 um; z-step size of 0.3 um). One or two ROIs per slide were collected in each nucleus (rostral to caudal extent of the ventral midbrain, n=5 cases, 6 slides/case, 6 ROI/ case) for each experimental case. These confocal parameters for identifying fluorescently labeled contacts have been validated using structured illumination microscopy and electrophysiology in thick sections (Fogarty et al., 2013). Imaris software 9.6/10 (Bitplane) was then employed to quantify interactions between tracer labeled boutons and TH- or GAD1-IR cell bodies to define axonal ‘contacts’ on post-synaptic cell bodies/proximal dendrites as previously reported (Kelly et al., 2021). In the 3D module, axonal terminals were analyzed using ‘spots’ rendering through the Z stack for the level at which they appeared brightest. In the “slice view”, fiber thickness was determined (1.75 um) for all cases.

Following a series of interactive histogram adjustments based on voxel size, we accurately detected as many possible boutons a possible without creating artifact. In the 3D module, TH- and GAD1-IR cell labeling was isolated using the “surface” rendering option. A total of 6 images per ROI per case were analyzed with approximately 10 cells randomly selected cells isolated per image, giving an equal number of TH or GAD1-labeled cells per case. Each ROI consisted of a 5x4 60X tile resulting in (512)^2^ um. Tracer/cell interactions for cells in each case were calculated using the “shortest distance” module. Based on initial fiber thickness measurement, shortest distance parameters that were greater than >0.75 um from the post-synaptic cell surface were considered a partial overlap of cell/fibers based on previous reports (Wouterlood et al., 2007; Kelly et al., 2021) and counted as a contact. Each TH+ or GAD1+ cell body containing at least one tracer-IR contact was manually checked in 3D, and manual counts of contacts were performed for each labeled cell. Results were expressed as the proportion of TH- or GAD1-positive cells receiving at least one contact per counted per total neurons in each ROI.

#### Electron Microscopy: Tracer contacts on TH and GAD1-IR distal dendrites

Images were captured on a Hitachi 7650 Transmission Electron Microscope using a Gatan 11-megapixel Erlangshen digital camera and Digital Micrograph software. TIFF images were later exported into Adobe Photoshop and Adobe Illustrator (v2022) and adjusted for brightness and contrast in preparation for analysis. EM analysis was done on thin sections collected from the surfaces of tracer-TH and tracer-GAD1 double labeled sections after selecting and excising ROIs of anterogradely labeled terminals at the light microscopic level in two animals. For the PBP and the A8 samples, all immunoreactive terminals were photographed (30K magnification) from samples containing detectable peroxidase and gold-silver labeling. Analysis was limited to sections where both peroxidase and gold-silver deposits were seen, to prevent false negatives due to uneven penetration of the reagents. Images in 1000 um^2^ of tissue were collected in PBP and in A8 regions for each animal (where fibers were most densely concentrated) in each double labeled set of FR/TH and FR/GAD1 labeled sections (total 2000 um^2^ for each tracer/transmitter per animal= total of 4000 um^2^ per animal). Terminals were then assessed for synaptic profiles and their type, and recorded. The number of axon terminals in each 1000 um^2^ region was recorded using the criteria below. Finally, FR+ terminals with synaptic specializations were assessed with respect to contacts onto TH-positive dendritic profiles or spines in the PBP and A8. This analysis was repeated for FR/GAD1 cases. We focused on labeled ATs on TH-IR and GAD1-IR dendrites or spines in each double-labeling experiment.

While ATs also contacted non-TH (or non-GAD1) dendrites in each compartment, their identity could not be inferred given the heterogeneity of the post-synaptic cells in each region. We refer to dendrites and spines together, as ‘dendrites’.

The following classifications were applied to characterize labeled axons, axon terminals, and dendritic elements throughout the neuropil, which were the main structures of interest.

##### Axon terminals, axon shafts

Immunoreactive processes were defined as axon terminals if they measured at least 0.2 um in diameter (Pickel et al., 1995), and were distinguished from other subcellular profiles based primarily on the presence of small synaptic vesicles and mitochondria. Axon terminals often had synaptic profiles with neighboring elements. In contrast to terminals, axon shafts appeared round when viewed in the transverse plane (i.e. axon bundles), and elongated when cut longitudinally.

##### Dendritic shafts

Dendritic shafts cut longitudinally were recognized by their irregular contours, elongated mitochondria in parallel with their central axis, frequent protuberances (spines, filopodia, small branches), and synaptic contacts with axon terminals. When cut transversally, dendritic shafts were identified by their rounded morphology, frequent occurrence of mitochondria and microtubules, and were distinguished from unmyelinated axons by their larger diameter. Dendrites appear more electron-lucent than axon shafts.

##### Dendritic Spines

Dendritic spines cut longitudinally often protruded from dendritic shafts, displayed rounded morphologies and were free of mitochondria. Spines were characterized primarily by the presence of electron-dense accumulations (postsynaptic densities) at synaptic contact sites with axon terminals.

##### Astrocytes

Protoplastic astrocytes were recognized as electron-lucent structures seen to encase and wrap around other neuropil structures. As a result, astrocytes maintained irregular and angular shapes, distinguishing them from other neuronal profiles having a characteristic rounded shape.

##### Synapse classification

‘Synapses’ were classified as tracer-IR axon terminals that made either a symmetrical or asymmetrical synaptic profile onto a post-synaptic element (usually a dendrite). A synaptic terminal was considered tracer+ if it contained either punctate DAB deposits or immunoreactive dense core vesicles, or both. All synapses were identified by thickened parallel membranes with a widened cleft. Asymmetric synapses had thickened post-synaptic densities (PSDs), generally greater or equal to 40nm (Peters et al., 1991; Dosemeci et al., 2016; Root et al., 2018). Symmetric synapses had thin post-synaptic densities with a considerably narrower synaptic cleft than asymmetric synapses (about 12 nm wide; <u>Peters et</u> <u>al., 1991</u>)

### Statistical analyses

Statistical analyses were performed using GraphPad Prism software (version 10). *Confocal data*: A two tailed Mann-Whitney test was used to compare the difference between medians of the proportions of CeN originating contacts onto TH+ or GAD+ soma/proximal dendrites cells in the ventral midbrain (**Fig. 4E-F**). A two-way ANOVA with Tukey’s multiple comparisons tests were used to investigate the frequency of contacts per cell type across midbrain regions as well as the comparison between ≤2 and >2 contact groups across regions and cell type (**Fig. 4G-H**). *Electron Microscopy data:* Two-tailed Chi-Square tests were used to compare the difference between total symmetric vs asymmetric synapses (**Fig. 6A**) and differences between synapse type across midbrain region (Fig. 6B) and dendritic cell type (**Fig. 6C-E**). Sex differences were considered in both the confocal and EM data using a two-tailed Mann Whitney test and were not significant from one another. p<0.05 was deemed significant. Error bars indicate SEM.

## Results

### Organization of DA/GABA neurons in the primate midbrain

The DA neurons of the rostral and caudal midbrain for one young male animal were qualitatively assessed for TH and GAD1 mRNA expression in specific subregions, and were compared to CaBP protein distribution in neighboring sections (**Fig 2A-A’ and B-B’**). As can be seen, TH- mRNA positive cells were found in both the CaBP-IR A10 and A8, and also densely clustered in the CaBP-negative A9. Somewhat surprisinging, there were relatively few TH-positive cells in the rostral midline of the A10, known as the rostrolinear nucleus. Throughout the midbrain, TH (red) and GAD1-positive neurons (green) formed largely separate populations. GAD1 mRNA was found in many neurons interspersed among all DA subregions, and in the SNr. A cluster of GAD mRNA cells were localized to the rostromedial tegmental nucleus (RMTg) and the interpeduncular nucleus (IP), which did not have TH-positive cells.

**Figure 2.**
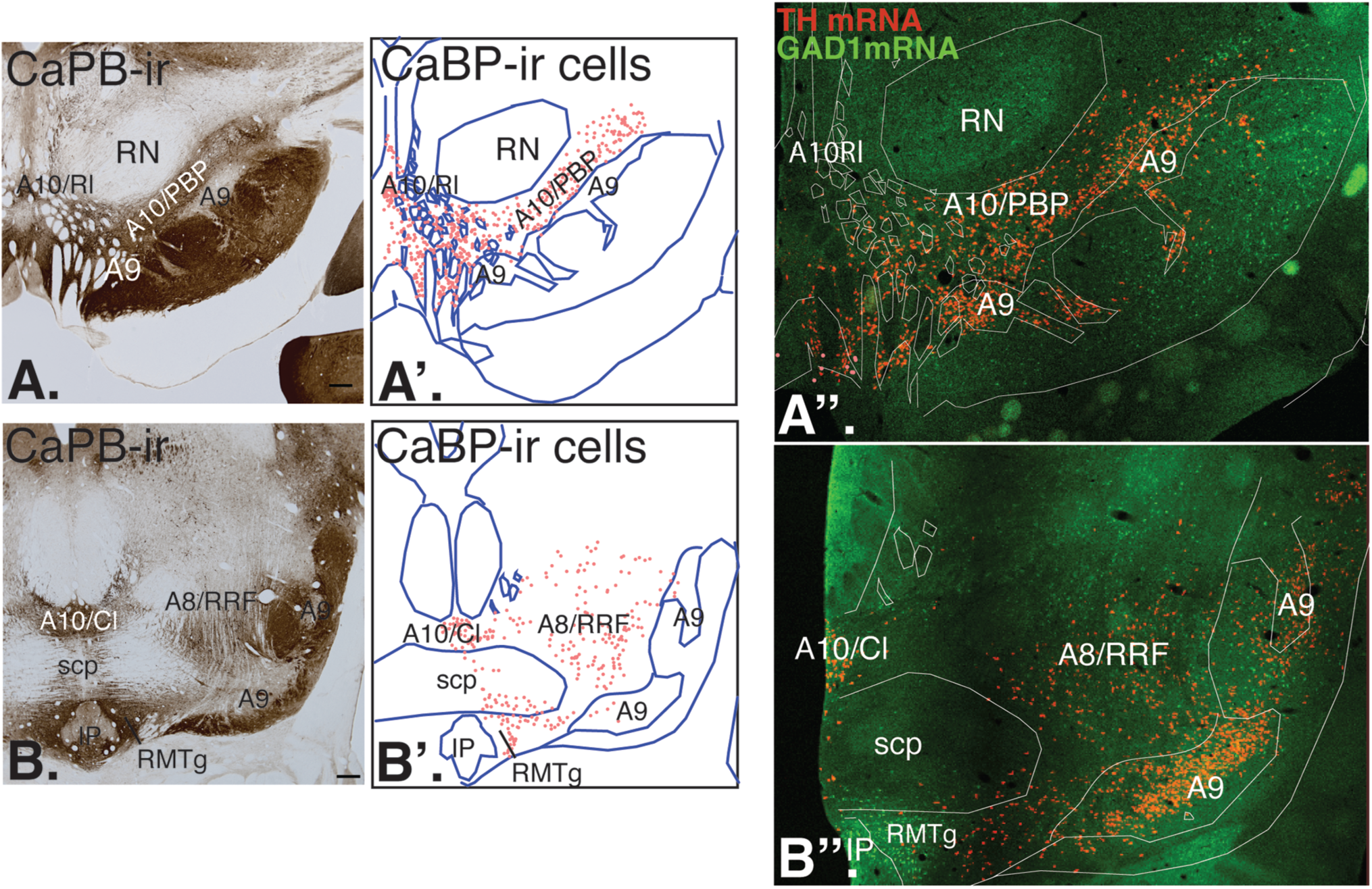
A-B. Low power photomicrographs of CaBP-IR in the rostral (A) and caudal (B) midbrain. A’-B’. CABP-labeled cell bodies are charted and distinguish the A10 and A8 neurons. The A9 neurons lack CaBP. The dense CaBP-IR in the SNr is in neuropil. A’’-B’’. Adjacent sections processed for THmRNA and GAD1mRNA, and aligned with CaBP-IR sections for subregional delineation. *A8/RRF, A8 retrorubral field; A10Cl, A10 caudolinear subnucleus; A10Rl, A10 rostrolinear nucleus; IP, interpeduncular nucleus; RMTg, rostromedial tegmental nucleus; RN, red nucleus;*

### Amygdala-nigral path

The distribution of tracer labeled fibers in the midbrain was similar following all 5 injections in the CeN. All injections resulted in labeled fibers over the dorsal tier (A10 and A8) neurons, with slight encroachment in some cases onto the dorsal A9 neurons (**Fig. 3**). In general, the distribution of labeled fibers in the ventral midbrain was very similar for all injection sites, regardless of the relative placement in the CeM versus CeL. All cases had labeled terminals overlapping with most densely over the PBP and the A8 (RRF) neuronal groups, while the midline VTA subnuclei were devoid tracer-positive fibers, as previously reported (Fudge et al., 2017). The inclusion of females in the present work did not reveal any clear differences in this pattern, including in assessment of terminal characteristics at the EM level (see below).

**Figure 3.**
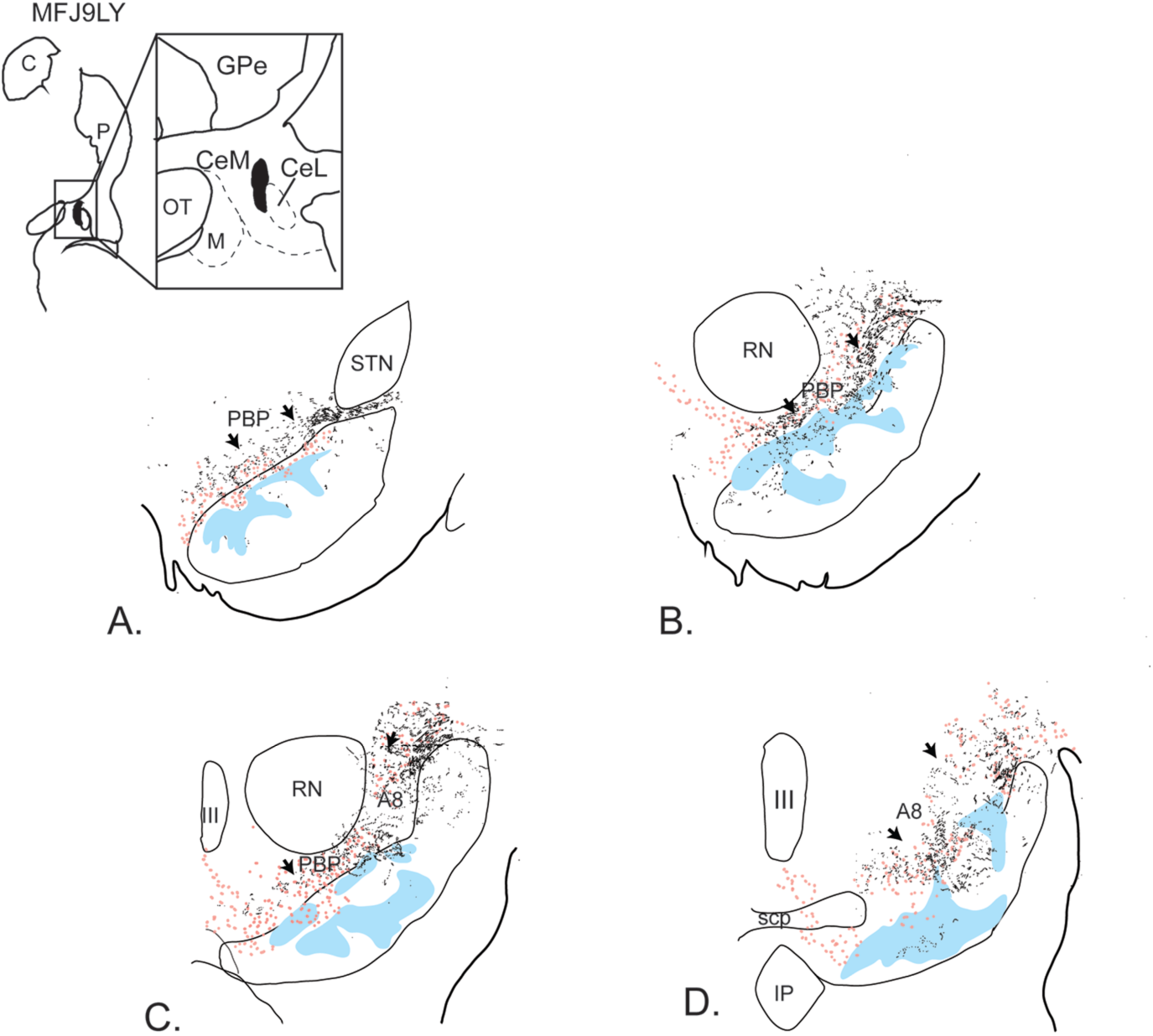
Representative map of a CeN injection site in case MFJ9LY, which is centered in the CeM. A-D. Rostrocaudal sections with anterogradely labeled fibers (black) hand-charted and superimposed on adjacent sections labeled and mapped for CaBP-IR cells (pink). Note that the labeled fibers avoid the midline at all levels and are distributed in the PBP and A8 (arrows), with slight encroachment on the A9. The CaBP-negative A9 region is depicted in blue. *III, third nerve; CeM, medial subdivision central nucleus, CeL, lateral subdivision of central nucleus, GPe, globus pallidus, pars externa; IP, interpeduncular nucleus; PBP, pigmented parabrachial nucleus (A10); RN, red nucleus; scp, decussation of the superior cerebellar peduncle; STN, subthalamic nucleus*.

### CeN afferent contacts onto DA and GABAergic soma and proximal dendrites at the confocal level (Fig. 4)

Using a mask for cell soma and proximal dendrites (**Fig. 4A-D**), we assessed 594 TH-positive neurons and 513 GAD1-positive neurons overall for potential terminal contacts (TH: PBP=311, A8=283; GAD1: PBP=251, A8= 262). Of these, 49% of TH-positive neurons (294/594) were contacted by at least one terminal; 52% of GAD1-positive neurons were contacted by at least on terminal (269/513). Thus, there was an equally likely chance of a terminal contact in each population. Moreover, this relationship was similar in the PBP and A8 for each cell type (TH: PBP 170/311, or 55%; A8: 124/283, or 44%; GAD1: PBP 135/251, or 54%; A8 134/262, or 51%) (**Fig. 4 E, F**, two tailed Mann-Whitney test, p=0.89 and 0.48 respectively).

**Figure 4.**
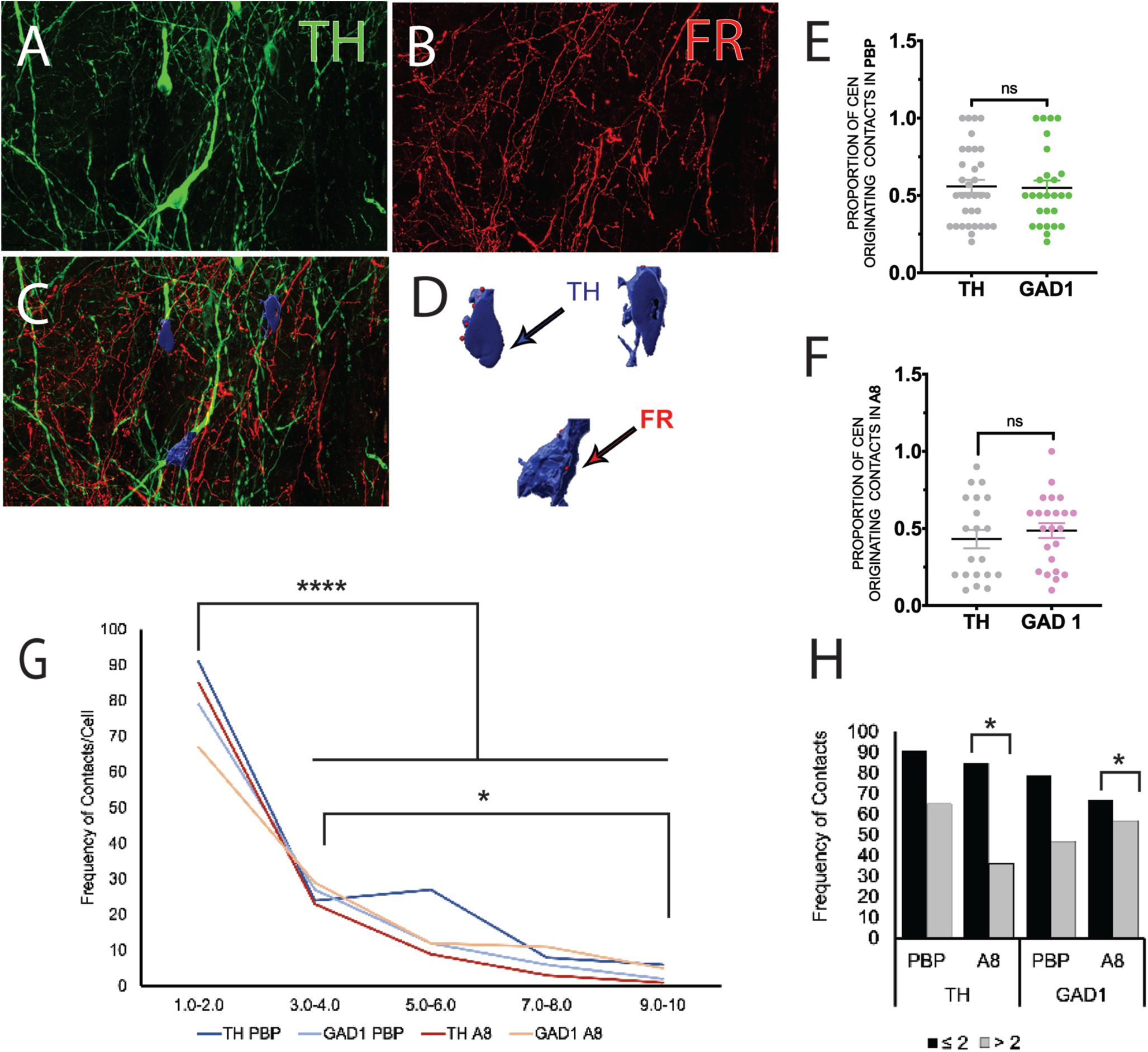
CeN afferent contacts on soma/proximal dendrites on TH- and GAD1-IR neurons. A-D. Method of confocal analysis. In section labeled for TH-IR neurons (green, A) and FR (tracer)-positive fibers (red, B), a mask of soma/proximal dendrites (blue) is created by hand (C). ‘Spots’ (FR-labeled contacts) that are less than 0.75 µm from the surface are counted (D). This was also done in GAD1- and tracer-labeled compartments in the same animal. E-F. The proportion of CeN tracer contacts on TH versus GAD-67-IR cell soma/proximal dendrites is strikingly similar, and is not different in the PBP versus A8. 1 dot= the proportion of 10 cells (1 ROI) with at least one contact. G. The frequency distribution of contacts onto TH and GAD1-IR cells in the PBP (blues) and A8 (reds). The majority of all cells in both regions receive 1 or 2 contacts/cell. H. Binning frequency data into the proportion of neurons receiving ≤ 2 or >2 contacts in each region indicated statistical differences between the bins, except for TH-PBP neurons.

The number of axonal contacts per neuron ranged from 1-10, with the majority of post-synaptic neurons received 1-2 contacts, regardless of cell type or region (**Fig. 4G**, p<0.000, F (4,12)=111.01, bin variation, two-way ANOVA; p<0.0001, 1-2 contact bin comp to all others, Tukey’s multiple comparisons test). The likelihood of a neuron receiving 1-2 contacts versus >2 contacts was statistically greater for both TH-IR cells in the A8, and for all GAD1-positive cells. The exception was for TH-IR cells in the PBP, in which there which there was an increased frequency in the 4-6 contact bin (**Fig. 4H, F**(3,3)=49.08, p=0.0048, regional variation, two-way ANOVA; Tukey’s multiple comparisons test, see **Fig 4G**) . On average, there were 3.9 contacts/TH+ cell, and 3.5 contacts/GAD1+cell in the PBP, and 2.4 average contacts/TH+ cells, and 3.8 average contacts/GAD1 neuron in the A8. Sample sizes were too low to assess sex differences, however, the counts in each animal (one male, one female) were similar. In sum, neuronal phenotype did not predict the chance of a CeN afferent contact, and the vast majority of neurons received 1-2 contacts on the soma and/or proximal dendrite.

### CeN afferent contacts onto TH- and GAD1-IR small dendrites

Electron microscopic examination of the PBP and A8 regions of interest indicated that reaction product for anterograde tracer was localized in unmyelinated axon shafts and axon terminals. We focused on axon terminals (ATs) to quantify their frequency and types of synaptic specializations. Reaction product was found in grainy depositions and also sequestered in dense core vesicles in ATs **(Fig. 5A-D**, pink structures). Of the total number of 459 ATs counted (226 in MFJ55 female, and 233 in MN1 male, **Table 2**), all had one or more synaptic specializations except for 27, which we termed ‘appositions’ (5.9% of all ATs). It is possible that appositions in nearby planes of the section did contain a synaptic specialization. This could not be assessed in our material. An occasional astrocytic process was seen interposed between pre-synaptic terminals (pink) and dendritic processes (blue) (**Fig. 5B, asterisks**), where they abutted synaptic contacts, or so-called ‘active zones’. Astrocytic processes surrounding synaptic contacts reports have been previously reported in studies of amygdala afferents to other catecholamine cell groups in rodents (Pickel et al., 1995; Pickel et al., 1996). Quantitative analysis of the combined cases showed that of the total of 459 anterogradely labeled ATs, 232 (50.5 %) were in the PBP and 227 (49.0%) of which were in the A8 (J1FR PBP= 119, J55FR PBP=113; J1FR A8=114, J55FR A8=113). The majority of ATs had synaptic specializations (∼90-95%), which were further analyzed.

**Figure 5.**
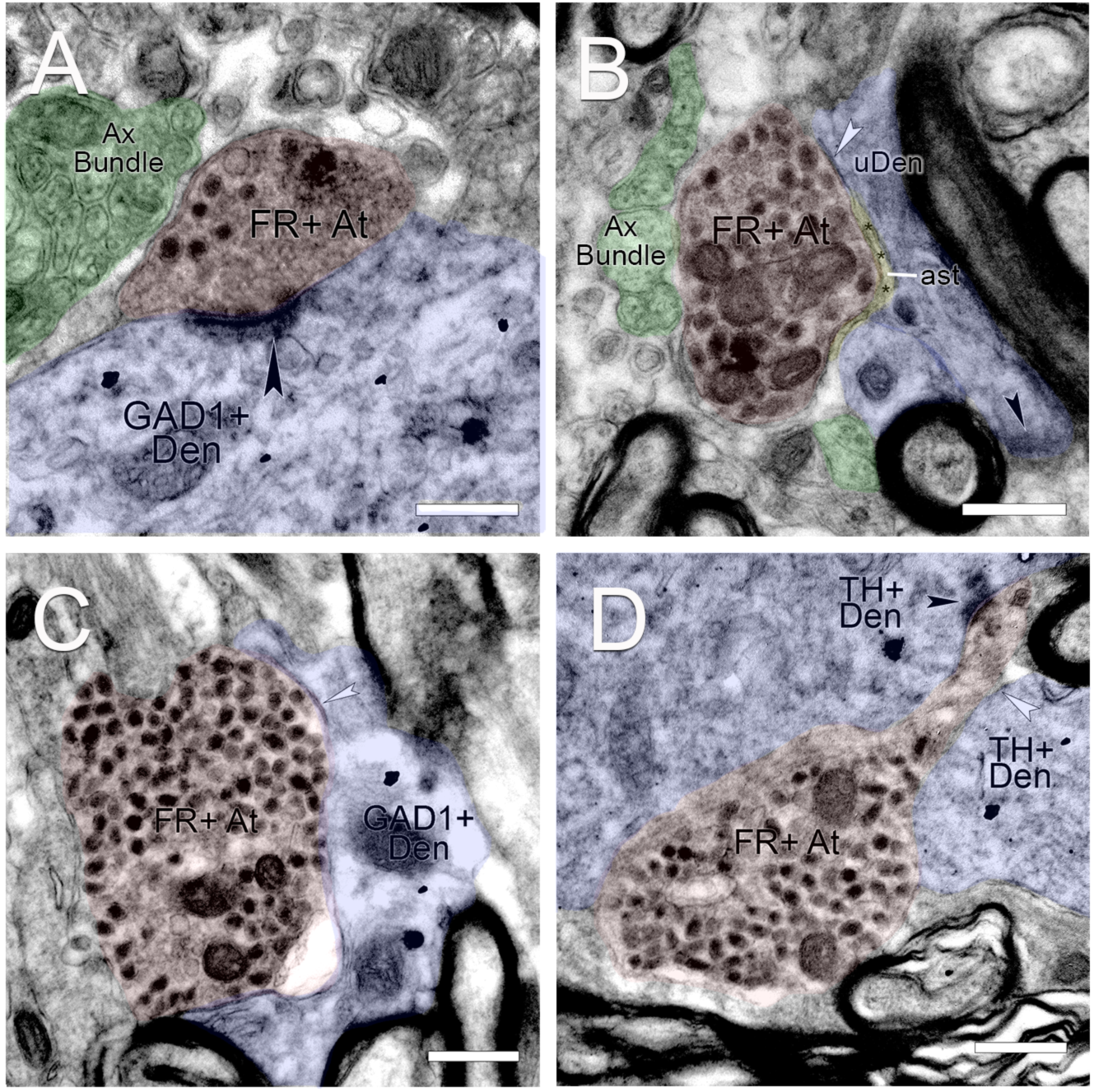
Representative examples of dual immunoperoxidase electron microscopy. A. Anterograde tracer labeling following CeN injection in characteristic densely filled vesicles containing tracer in the midbrain (pink FR+At). Reaction produce is in dense core vesicles and also seen as particulate deposits. Axon shafts in an axon bundle (green). GAD1-Gold labeling of a neighboring dendrite (purple) receiving an asymmetric contact. B. FR+ At (pink) making an symmetric (putative inhibitory) contact (white arrowhead) onto an unlabeled element (uDen, purple). Note-astrocytic sheath (ast, asterisks) surrounding terminal. C. FR+At terminal (pink) making a symmetric-type contact (white arrowhead) onto a GAD1+ Den (purple). D. FR+At terminal (pink) bordered by two TH+ Den (purple), making an asymmetric profile (black arrowhead) onto one TH+ element, and symmetric-type synapse onto the other (white arrowhead). Scale bar= 500nm.

### CeN afferents form symmetric and asymmetric-type synapses onto dendrites

The majority of tracer-labeled ATs contacted dendritic profiles, including spines, in our two series through the midbrain (tracer/TH and tracer/GAD) of each animal. A total of 534 synaptic profiles were seen on labeled ATs, and the vast majority were onto dendritic shafts or spines (26 synaptic contacts onto unidentifiable elements, 5.7%). Synaptic profiles were predominantly symmetric (putative inhibitory) (n=303, 57%) profiles, with asymmetric-type (putative excitatory) profiles comprised 43% (n=231) of the total (I/E ratio 1.3 overall) (**Fig. 6A**, **Table 2**). There were no differences in this relationship between the PBP and A8 portions of the terminal field (**Fig. 6B**). Occasionally, both synaptic types were found on the same AT (e.g. **Fig. 5D**).

**Fig. 6.**
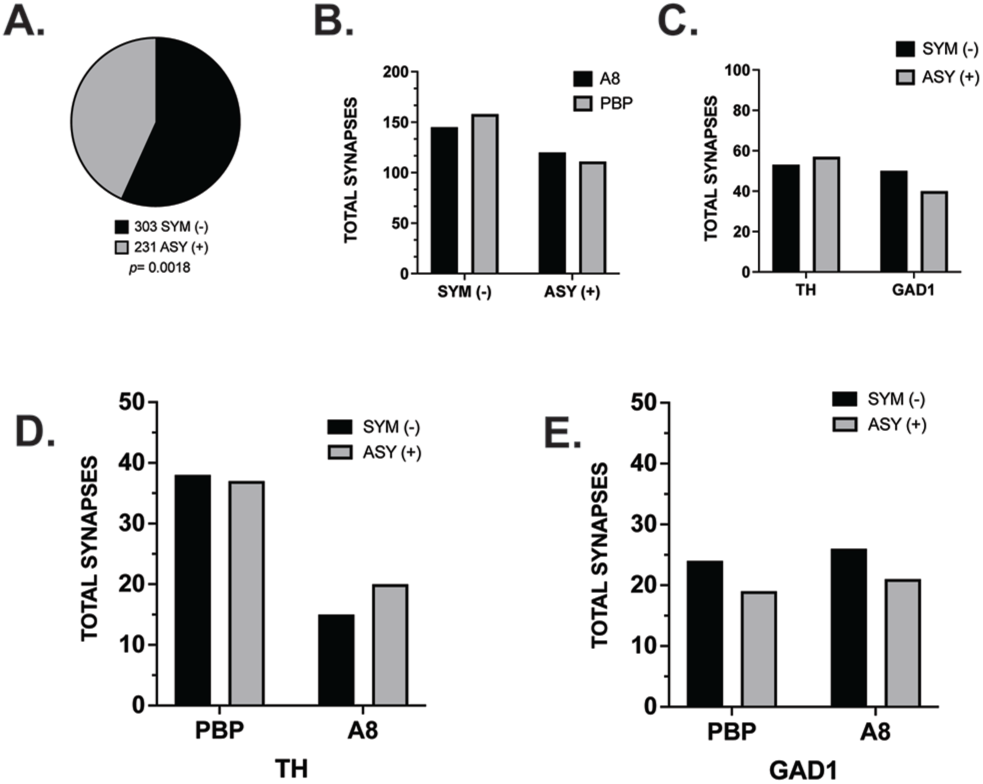
CeN afferent contacts at the EM level. A. Comparison of all symmetric (putative inhibitory) versus symmetric (putative excitatory) profiles on all labeled axon terminals. B. Symmetric versus asymmetric terminals in the PBP and A8 are similar. C. Total symmetric (black) and asymmetric (gray)-type contacts onto GAD1 and TH-positive elements. No significant differences between synapse type onto either GAD1 or TH-IR dendrites, or between regions. (two-tailed Chi-Square, p=0.3). D. The distribution of asymmetric and asymmetric-type synapses onto TH labeled dendrites in the PBP and A8 (two-tailed Chi-Square, p=0.45). E. The distribution of asymmetric and asymmetric-type synapses onto GAD1 labeled dendrites in the PBP and A8 (two-tailed Chi-Square, p=0.96).

### Labeled axon terminals on TH- and GAD-positive dendrites

We collected FR/TH and FR/GAD1 samples of equal area for each region, in each animal. Tracer labeled ATs made a contact on labeled post-synaptic dendrite and/or spine--versus a non-labeled element--approximately 39% of the time regardless of compartment (FR-TH, 110/284, 38.7%; FR-GAD1, 98/250, 39.2%, **Table 3**). With respect to contacts onto TH-positive dendrites, 48% (53/110) were symmetric and 52% (57/110) were asymmetric (black arrowheads)(**Fig. 6C**, two-tailed Chi-Square, p=0.3, **Table 3**). In tracer and GAD1-IR adjacent sections, 59% of synaptic profiles on tracer-labeled ATs that contacted GAD1-positive dendrites were symmetric (58/98), and 41% were asymmetric (40/98) (**Fig 6C**, two-tailed Chi-Square, p=0.3).

Comparing asymmetric versus symmetric contacts associated with TH-positive dendrites, both synaptic types were higher in the PBP compared to the A8 region. However, there was no difference in the relative number of asymmetric versus symmetric type profiles in either region (Two-tailed Chi-Square, p=0.45) (**Fig. 6D**). In FR/GAD1 labeled sections, the total number of synapses between the PBP and A8 were similar, as was the number of asymmetric versus symmetric synapse types within each region (Two-tailed Chi-Square, p=0.96) (**Fig. 6E**).

## Discussion

This study characterized the primate amygdalonigral path at a cellular/synaptic level in the nonhuman primate. Central nucleus inputs to the ventral midbrain have been implicated in a range of behaviors, including orienting to salient sues(Lee et al., 2006), and reinforcing both rewarded and defensive behaviors(Steinberg et al., 2020). Despite encompassing a broad continuum of ventral midbrain, we found striking uniformity in the basic synaptic characteristics of CeN afferent terminals and their propensity to contact either DA or GABAergic cells.

Although the PBP and A8 regions are large, with cellular and connectional heterogeneity, terminals from the CeN had an almost equal chance of contacting DA versus GABA neurons in each region. With few exceptions, CeN terminals also followed similar patterns in terms of contact frequency onto each dendritic compartment, for both DA and GABA-labeled structures, in the PBP and A8.

### DA and GABAergic neurons across DA subregions

The classic DA subregions were examined with respect to TH and GAD1 mRNA distribution, to illustrate the distribution of putative DA and GABAergic cells. In general, there was a strong separation of TH- and GAD1R-mRNA signal throughout ventral midbrain, with little colocalization. GABA is known to be loaded into DA neurons by the vesicular monoamine transporter (VMAT) for co-release at terminals(Zych and Ford, 2022), but its production exists outside the DA neuron (Tritsch et al., 2014). We found GAD1 mRNA positive cells were evenly distributed across the ventral midbrain, while TH mRNA labeled neurons were generally denser and had higher variability. These qualitative results support our recent quantitative stereologic results in same age animals using TH- and GAD1-protein expression (Kelly et al., 2022). In that study, TH-IR neurons were more abundant in the PBP compared to the A8, while GAD1-IR cells were more evenly distributed across all regions. In the PBP and A8, the ratio of TH-GAD1 labeled neurons in the PBP was 5:1, while in the A8 the ratio was 1:1. Thus, there are considerable differences in the ‘weighting’ of DA and GABA cell type across these midbrain regions that receive CeN afferents.

### CeN inputs project with equal probability onto DA and GABAergic neurons

CeN injections all had a similar projection to the midbrain, with labeled fibers concentrated mainly over the PBP (or ‘lateral’ VTA) and A8, as previously reported(Fudge and Haber, 2000; Fudge et al., 2017). Relatively light patches of labeled fibers encroached on the dorsal A9 and into the midline VTA (A10) in some injections.

Examining and quantifying terminals in the PBP and A8 under higher magnification, we found that the propensity of CeN labeled terminals to contact either DA or GABAergic cell type was approximately equal, regardless of positions at proximal versus distal regions of the dendritic arbor. A reason for examining afferent synapses onto cell soma and their proximal dendrites is that these contacts physiologically privleged, having an outside effect on inducing action potentials (Hausser et al., 1995; Seutin and Engel, 2010; Moubarak et al., 2022). On DA neurons, the axon hillock is often found on large diameter proximal dendrites close to the soma(Juraska et al., 1977; Grace and Bunney, 1983). Here, somatodendritic morphology is plays a causative role in action potential initiation and duration(Hausser et al., 1995; Meza et al., 2018; Moubarak et al., 2022). In GABAergic neurons, action potentials almost always begin at the soma where the axon originates (Hausser et al., 1995). Because the soma and thick proximal dendritic structures are less likely to be captured at the resolution of EM, confocal methods were used. Interestingly, on soma and proximal dendrites DA and GABAergic neurons receiving a contact, most received 1-2 contacts, regardless of transmitter phenotype. Smaller distal dendrites of the neuropil, comprising the vast majority of the dendritic arbor, were also equally likely to receive a contact regardless of transmitter type. In equal areas of neuropil, tracer-positive terminals made synaptic contacts onto TH- and GAD-labeled small dendrites 39% of the time (compared to contacts on no-labeled dendrites).

Interestingly, this work is congruent with recent rodent studies using retrograde viral techniques to examine afferents to specific neuronal phenotypes midbrain(Beier et al., 2015; Faget et al., 2016; Beier et al., 2019). Despite different methodologies and species, afferent inputs detected in these studies (which ‘seed’ specific cell populations and use viral tracing) also fail to show afferent specificity or preference for specific neuron transmitter types. There present study shows this on a single afferent path in monkey, across a large spatial target region, and at several levels of the dendritic compartment.

### Unbiased afferent inputs to shifting DA:GABA ensembles

The amygdalonigral path is only one of myriad afferent paths that modulate DA dynamics, but is instructive in understanding how afferent paths influence DA subregion outputs generally. The present results are strikingly similar to previous studies, as reviewed above, finding little bias in afferent contacts for either neuron type. The strong uniformity in CeN contacts on DA and GABA neurons across a broad span of the PBP-A8 raises the possibility that post-synaptic cellular organization critically determines output channels.

While much work has focused on the molecular diversity of the DA system, and circuit-specific associations (Poulin et al., 2018; Beier et al., 2019; Derdeyn et al., 2021; Azcorra et al., 2023), much less is known about GABAergic neurons, including their relative densities among DA neurons, and how GABA shift across the midbrain affect DA output. GABAergic neurons serve as interneurons and long-range projections(Carr and Sesack, 2000; Omelchenko and Sesack, 2009; Dobi et al., 2010; Taylor et al., 2014; Breton et al., 2019). In primates, we recently found that their relative densities in relation to local DA cells (but not each other) shifts strikingly along the mediolateral and rostrocaudal plane(Kelly et al., 2022). For example, we found five-fold changes in DA neuron-to-GABA neuron densities (and morphologic diversity) are found in the PBP compared to the A8. Within the CeN projection to this entire region, spanning the PBP and contiguous A8, rostrocaudally, each DA and GABA neuron has a ∼40-50% chance of receiving a contact despite the large rostrocaudal and mediolateral expanse of the region. However, because the ratio of DA:GABAergic neurons shifts across from the PBP to A8, how amygdala contacts influence DA versus GABA outputs also shifts regionally **(Fig. 7).** Thus, while axons may contact a fixed proportion of DA or GABA cells, creating a seemingly ‘unbiased’ set of inputs, the fact that the relative densities of post-synaptic neuron phenotypes change their balance with respect to one another is an important context to consider in understanding the input effects on outputs in DA subregions.

**Fig. 7.**
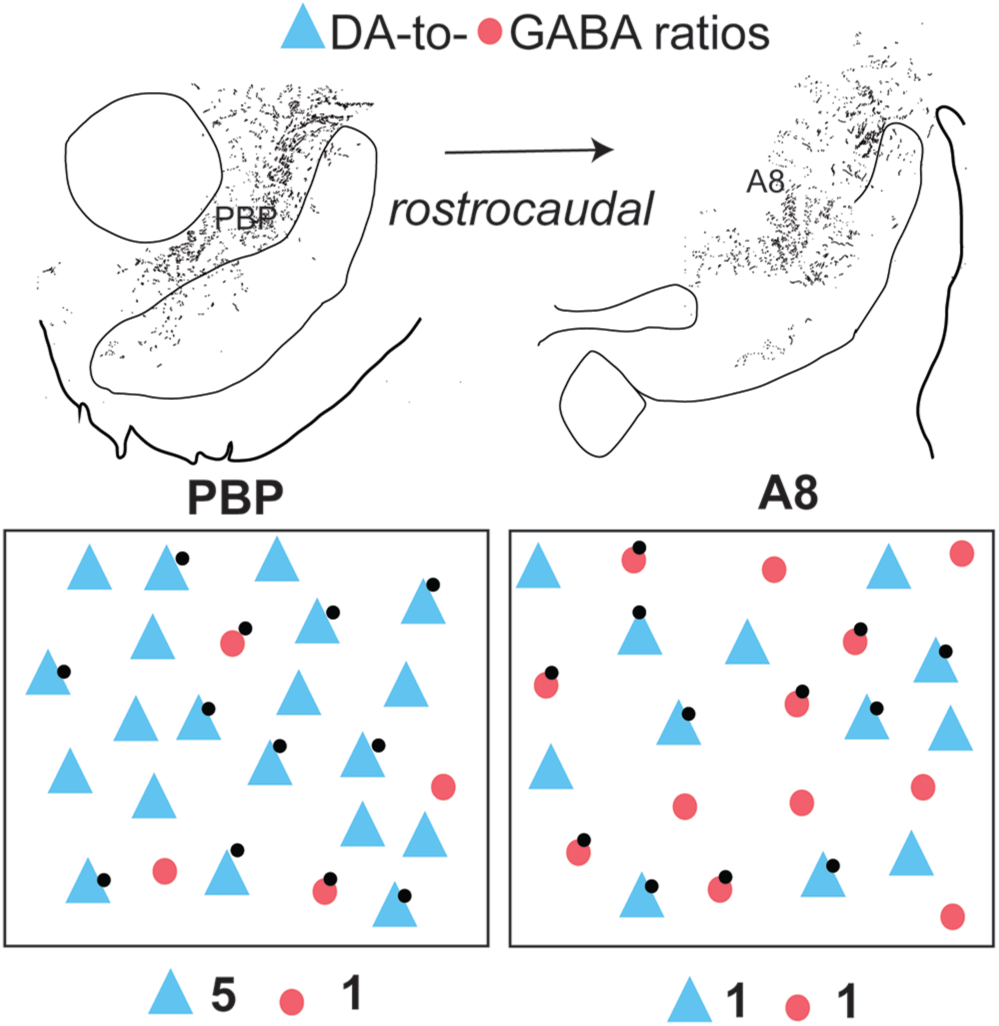
CeN contacts in the context of shifting DA:GABA ratios in the PBP and A8. While each phenotype has 40-50% change of receiving a contact (versus not), shifting ratios of DA:GABA (blue:red) neurons mean that each subregion is physiologically unique.

Rules of DA neuron physiology and their respective output streams correlate with their position along the mediolateral and rostrocaudal axis of the primate ventral midbrain (Mirenowicz and Schultz, 1996; Matsumoto and Hikosaka, 2009). In primates, ‘reward-prediction error’ coding neurons tend to localize in the medial VTA, while ‘salience’ coding responses (to stimuli that are novel or threatening, and involved in ‘orienting’ and threat responses) are located laterally and caudally. These topologic relationships are generally consistent with rodent data (e.g. (Cohen et al., 2012; Menegas et al., 2018; Moaddab and McDannald, 2021)). However, in primates the expanded PBP and A8, which are outside the midline VTA, house multiple output streams, including efferents to the central and posterior striatum, the extended amygdala and amygdala. PBP and A8 efferents are widespread and overlapping in the nonhuman primate. Both the PBP and contiguous A8 are a major source of projections throughout the amygdala (Cho and Fudge, 2010) and much of the prefrontal cortex (Gaspar et al., 1992; Williams and Goldman-Rakic, 1998). The PBP projects to broad regions of the primate limbic and associative striatum (Haber et al., 2000), while the A8 neurons innervate the caudal ventral striatum(Francois et al., 1999).

Paradoxically, then, inputs from the CeN which favor neither DA or GABA neurons when these data are normalized, are poised to differentially influence specific outputs to all these regions due to strong shifts in DA:GABA ratios across the region.

## ACKNOWLEDGEMENTS

We thank the URMC Electron Microscope Research Core and URMC Center for Advanced Microscopy and Nanoscopy Core (CALMN) for their expert guidance and assistance, Nanette Alcock (University of Rochester SMD) for help with histological preparations, and funding by the National Institutes for Mental Health (RO1MH115016, J.L.F.)

