## Supplementary material for "Amygdalo-nigral inputs target dopaminergic and GABAergic neurons in the primate: a view from dendrites and soma": Table 1

| **Animal ID and**  **tracer** | **Sex** | **Age (years)** | **mRNA** | **Tract tracing** | **Contacts on Soma/proximal dendrites: confocal** | **Contacts on small dendrites: EM** |
| --- | --- | --- | --- | --- | --- | --- |
| J01FR | Male (*m. nemestrina*) | 1.4 |  | XX | X | X |
| J09LY | Male (*m. fascicularis*) | 4.2 |  | XX | X |  |
| J38-- | Male (m. fascicularis) | 3.3 | X |  |  |  |
| J55FR | Female (*m. fascicularis*) | 2.9 |  | X | X | x |
| J56FR | Female (*m. fascicularis*) | 3.3 |  | X | X |  |
| J58FS | Female (*m. fascicularis*) | 3.6 |  | X | X |  |

Table 1. Animal characteristics and use.
