## Supplementary material for "Amygdalo-nigral inputs target dopaminergic and GABAergic neurons in the primate: a view from dendrites and soma": Table 2

| **Case** | **Labeled AT** | **All labeled ATs with**  **synaptic elements** | **All symmetric profiles (-)** | **All asymmetric profiles (+)** | **Symmetric to asymmetric ratio** |
| --- | --- | --- | --- | --- | --- |
| MFJ55FS (F) | 226 | 214 | 148 | 118 | 1.25 |
| MNJ1FS (M) | 233 | 218 | 155 | 113 | 1.37 |
| TOTALS | 459 | 432 | 303 | 231 | 1.31 |

Table 2. Afferent axon terminal (AT) characteristics in each animal.
