## Supplementary material for "Amygdalo-nigral inputs target dopaminergic and GABAergic neurons in the primate: a view from dendrites and soma": Table 3

| **FR-TH compartment** | | | | | |
| --- | --- | --- | --- | --- | --- |
|  | number (%) |  | number (%) |  | number (%) |
| All dendrites/spines |  | TH+ dendrite/spines |  | Unlabeled dendrites/spines |  |
| Symmetric | 162 (57%) | Symmetric type | 53 (48%) | Symmetric | 109 (63%) |
| Asymmetric | 122 (43%) | Asymmetric type | 57 (52%) | Asymmetric | 65 (37%) |
| Total | 284 |  | 110 |  | 174 |
| **FR-GAD1 compartment** | | | | | |
| All dendrites/spines |  | GAD1+ dendrite/spines |  | Unlabeled dendrites/spines |  |
| Symmetric | 141 (56%) | Symmetric type | 58 (59%) | Symmetric | 83 (55%) |
| Asymmetric | 109  (44%) | Asymmetric type | 40 (41%) | Asymmetric | 69 (45%) |
| Total | 250 |  | 98 |  | 152 |

Table 3. Synaptic contacts onto TH-positive and GAD1-positive

dendrites/spines versus unlabeled dendrites/spines.
